# The End of Aging Clocks: Training Foundation Models to Reason in Aging and Longevity

**DOI:** 10.64898/2026.03.28.714980

**Authors:** Alex Zhavoronkov, Vladimir Aladinskiy, Alex Aliper, Zulfat Miftakhutdinov, Mathieu Reymond, Vladimir Naumov, Diana Zagirova, Stefan Pushkov, Denis Sidorenko, Rim Shayakhmetov, Fedor Galkin

## Abstract

The aging clock paradigm has yielded dozens of specialist models that can estimate chronological age or mortality from virtually any biodata type. Yet each such model operates within a fixed modality, relies on a predetermined feature set, and produces limited biological interpretation. Here, we report Longevity-LLM v0.1, a Qwen3-14B model fine-tuned through supervised and reinforcement learning regimes on DNA methylation, proteomics, clinical biomarker, and RNA expression data. Longevity-LLM achieves high ranks in the recently announced Longevity Bench, including such tasks as cancer survival and RNA- or proteome-based age prediction. After reinforcement fine-tuning, the model achieved a 4.34-year MAE in epigenetic age prediction, surpassing the Horvath multi-tissue clock. In addition to age prediction, Longevity-LLM can carry out numerous other tasks, including proteomic profile generation, for which it significantly outperforms all frontier LLMs. These results demonstrate that a single modestly sized LLM can match or replace purpose-built aging clocks across data modalities.

This work constitutes an interim report from the initial sprint of our Multi-Modal AI Gym for Science (MMAI), an initiative dedicated to building foundation models for drug discovery and aging research.

## Introduction

Aging clocks have become indispensable tools in longevity research. Since the introduction of the first deep learning-based aging clocks in the mid-2010s, the field has produced specialized predictors for DNA methylation ^1–3^, gene expression ^4,5^, blood proteomics ^6–8^, clinical biomarkers ^9^, gut microbiome composition ^10^, and facial photography ^11^, among other data types ^12^. These models are now ubiquitously used in anti-aging experiments and trials.

Despite their utility, aging clocks share a set of fundamental constraints that greatly limit the scope of problems they can be applied to ^13,14^. Firstly, any clock is confined to a single biodata modality with a predetermined feature set. This limitation is a direct consequence of multimodal data scarcity accompanied by the inability of the utilized machine learning methods to operate with incomplete observations. Adding a new modality to a clock requires building, validating, and deploying an entirely new model. Moreover, such a multi-modal clock would only be applicable to similarly processed samples, effectively multiplying the cost of screening for each additional modality. From a viewpoint of interpretability, aging clocks also do not explain why a patient’s biological age exceeds their chronological age. Complementary analytic methods are required to interpret the aging signal they detect, such as differential gene expression, gene set enrichment, and graph-based approaches. Even with these tools, the interpretation is ultimately provided by the researchers themselves.

One solution to these problems is presented by AI agentic systems that combine a conversational model with external bioinformatic tools ^15,16^. While such systems can integrate data from multiple sources and present them in natural language, they do not resolve the underlying issues ^17,18^. The LLM in these architectures serves as an orchestrator routing queries to specialist tools and summarizing their outputs, rather than as a system that understands the biology itself. The specialist tools themselves remain frozen and carry over their biases and shortcomings to the orchestrator. This approach runs counter to our vision of a foundation AI that internalizes aging biology across modalities and deepens its understanding through iterative training.

In this work, we propose an alternative. Rather than equipping an LLM with aging clocks, we implemented aging clock distillation enabling an LLM to develop a multifaceted view of aging biology. We converted omics data and accumulated knowledge embedded in multiple aging clocks into structured traces for supervised and reinforcement fine-tuning (SFT, RFT) of a foundation model directly. The resulting system, Longevity-LLM v0.1, operates within a single 14-billion-parameter Qwen3 model and can process methylation, proteomic, transcriptomic, and clinical data without external dependencies. These advances enable creating a monolithic AI drug development pipeline, which would be able to independently handle all stages from target identification ^19,20^ to clinical trial design ^21^. While separate currently solutions exist to assist data analysis and decision making throughout a drug development program ^22^, we see a need in a system integrating the knowledge contained in single-purpose tools with generalist conversational LLMs, resulting in a pharmaceutical superintelligence, or a “prompt-to-drug” pipeline ^18^.

Here, we report the results of our initial ten-day development sprint. We evaluated the fine-tuned model against 15 frontier commercial LLMs on tasks from the Longevity Bench ^23^, compared its age prediction accuracy to established epigenetic and proteomic aging clocks, and assessed its capacity to generate biologically plausible proteomic profiles conditioned on age. These experiments represent the first application of our MMAI Gym for Science pipeline to the aging domain.

## Methods

### Training data

The training corpus comprised 766,640 examples (1.09 billion tokens) spanning 38 tasks across four primary data modalities, with an additional 191,804 examples reserved for holdout evaluation (**Supplementary File 1**). DNAm data constituted the largest modality (416,110 training prompts, 875 million tokens), including CpG beta value profiles for age regression, pairwise feature importance comparisons derived from published clock coefficients, and annotated CpG panels with genomic and pathway-level reasoning traces. Clinical biomarker data from NHANES ^24^ contributed 228,381 training examples across age prediction, mortality classification, and time-to-event tasks in multiple prompt formats. Transcriptomic data from GTEx Portal ^25^ comprised 98,510 training prompts for age prediction from gene expression profiles and pairwise tissue age comparisons. Proteomic data from Olink Explore 3072 plasma panels represented the smallest modality (7,807 training examples) with age regression, classification, pairwise comparison, and protein profile generation tasks. Data for proteomic prompt collections was obtained from ^26,27^ and the Immunobiology of Aging cohort described originally in ^28^ and accessed through the Allen Institute of Immunology web portal. Additional tasks in the corpus also included TCGA cancer survival comparisons (obtained through TCGA Research Network ^29^), multi-mutant lifespan prediction, and general aging biology tasks.

To improve Longevity-LLM’s generalization ability, prompt collections feature semantic variation, formatting options, synonym rotation, and automatically generated reasoning traces. Training and test splits were maintained at the subject level across all datasets to prevent data leakage. To explore the specific prompt schemas we used, see **Supplementary File 1**. The prompt collections featuring aging clock coefficients relied on public coefficients accessed via Biolearn ^30^.

### Model training

Longevity-LLM was derived from Qwen3-14B ^31^ through two sequential stages of full-parameter fine-tuning: supervised fine-tuning on a multitask mixture of aging-related prompt collections, followed by reinforcement fine-tuning on tasks with verifiable regression targets. No parameter-efficient adaptation methods were used in either stage.

#### Supervised fine-tuning (SFT)

The SFT stage consumed approximately 4.5 billion tokens over 3,000 optimization steps. The training mixture combined instruction-response pairs with examples containing intermediate reasoning traces, included jointly within a single training run to expose the model to a diversity of tasks. To mitigate overrepresentation of large datasets, sampling was performed uniformly across data sources and specific tasks.

Training used a 65,536-token context window with sequence packing, AdamW optimization (learning rate 1×10□□, cosine decay with 2% warmup, β□=0.9, β□=0.95, weight decay 0.1), FP16 precision, gradient checkpointing, and Flash Attention. The global batch size was 1,572,864 tokens per step. Training was conducted on 24 NVIDIA B200 GPUs.

#### Reinforcement fine-tuning (RFT)

RFT was initialized from the SFT checkpoint and applied Group Relative Policy Optimization to seven regression tasks where model outputs could be evaluated directly against known numerical targets (see **Supplementary File 1**). For each prompt, eight completions were sampled at temperature 1.4 and scored by a composite reward function. A Kullback-Leibler penalty (β=0.2) against the frozen SFT reference model prevented policy collapse.

The reward function comprised three components. A format reward required exactly one properly paired set of reasoning tags:

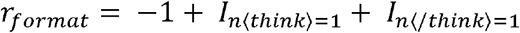

where *n*⟨think⟩ and *n*⟨/think⟩ count the occurrences of opening and closing tags, respectively. A thinking length reward incentivized substantive reasoning up to a saturation point:

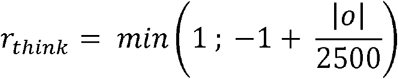

where |o| is the character count of reasoning content, saturating at 5,000 characters. A regression correctness reward evaluated prediction accuracy:

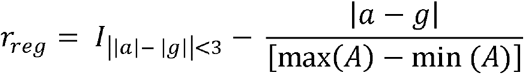

where ∣a∣ and ∣g∣ denote string lengths, and the second term is the normalized absolute error. The normalization denominator max(A) − min(A) is computed over the full range of target values in each task dataset, ensuring reward magnitudes are comparable across tasks despite differing native units (e.g. years or months). An additional penalty of −1.0 was applied for predictions outside task-defined validity bounds.

RFT used a 16,384-token context (2,048 prompt, 14,336 completion) without sequence packing, as each completion required independent reward evaluation. Optimization used AdamW (learning rate 6×10□□, constant schedule, β□=0.9, β□=0.99, weight decay 0.05) with gradient clipping at norm 0.05 and an effective batch size of 16 sequences. Training ran for 2,000 steps on 16 NVIDIA A100 80GB GPUs, with checkpoints saved every 500 steps and the final model selected by holdout evaluation performance.

### Evaluation metrics

Classification tasks were evaluated by accuracy (proportion of correct predictions). Regression tasks were evaluated by mean absolute error (MAE), Spearman rank correlation (ρ), and coefficient of determination (R^2^). The generative proteomic profile task was evaluated by the Jaccard index between the predicted and ground-truth protein sets. Confidence intervals for all metrics were computed by bootstrap resampling.

To compare the epigenetic age prediction performance of Longevity-LLM against the Horvath ^32^ multi-tissue clock, we applied a two-sample bootstrap test. Absolute prediction errors were independently resampled with replacement from both models (10,000 iterations), and the one-sided p-value was computed as the proportion of bootstrap replicates in which Longevity-LLM did not achieve a lower MAE than the Horvath clock. To keep the comparison fair, we used the study-level split strategy, as described in ^1^, which resulted in a test cohort with 1,436 samples not seen by either the Horvath clock or Longevity-LLM at training.

For proteomic clock comparison we used models with published weights: PAC ^7^, and Proteoclock ^6^. Test sets for this comparison was obtained from ^27^ and Immunobiology of Aging cohort ^28^, collected from the web portal of the Allen Institute for Immunology. 94 healthy samples from these cohorts were separated into a holdout set, not seen by the model in any form during fine-tuning stages.

All analyses were implemented in Python 3.11. Figures were prepared in Plotly with manual assembly in Adobe Illustrator.

## Results

### Performance in Longevity Bench

To place Longevity-LLM in the context of current commercial systems, we evaluated it against 15 frontier LLMs across seven Longevity Bench tasks spanning tasks in the domains of oncology prognosis, age and mortality prediction. Despite having only 14 billion parameters, our SFT model ranked first in four out of seven tasks (**Figure 1**). The base Qwen3-14B model, prior to fine-tuning, was unable to produce valid predictions on any of the tested tasks due to the inability to generate parseable outputs. The results displayed therefore have base model’s effectively void metrics removed.

**Figure 1.**
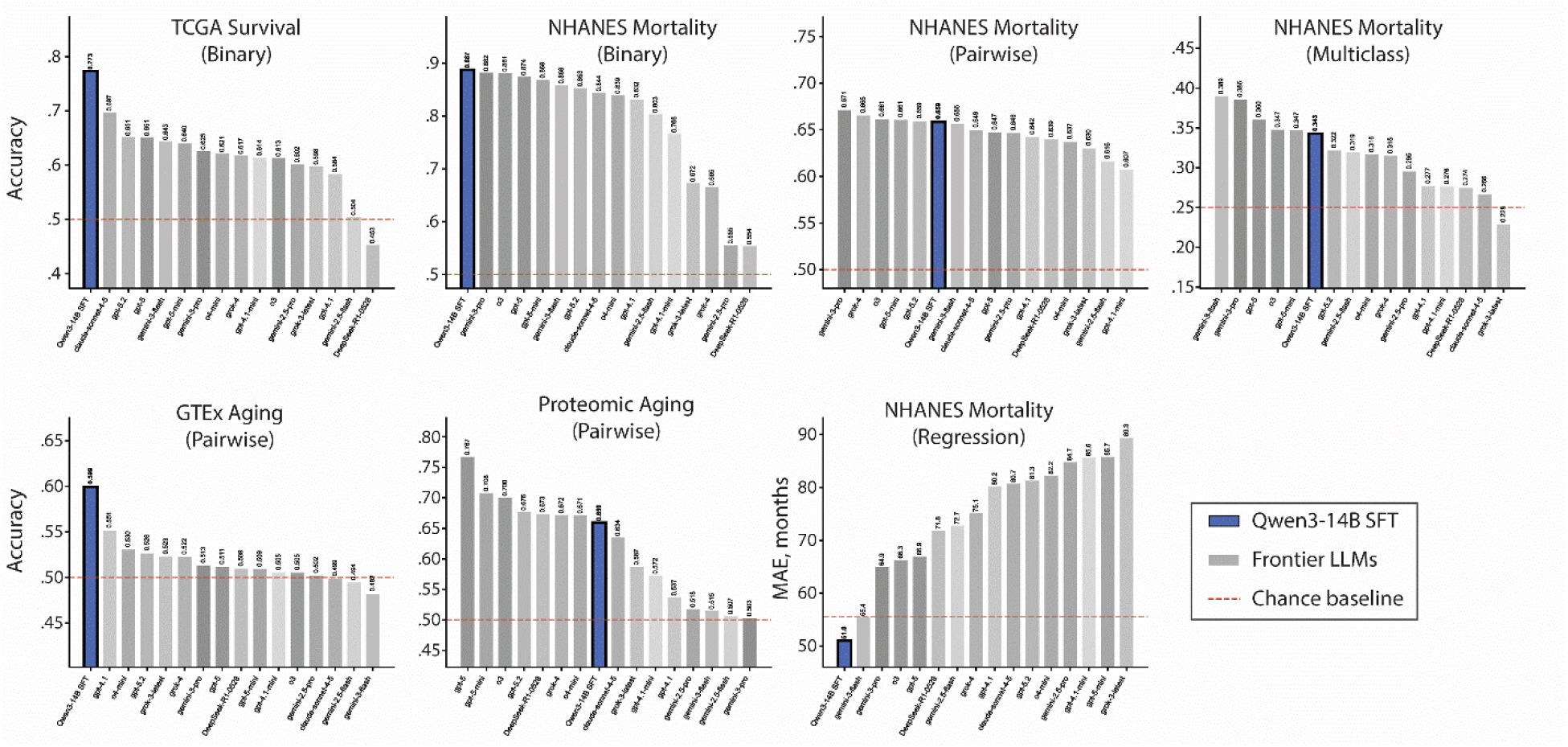
Longevity-LLM shows competitive performance in a range of Longevity Bench tasks. The tested tasks span the domains of transcriptomics, proteomics, and clinical record. Base Qwen-3 14B failed to produce valid outputs on all tasks.

The most pronounced advantage was observed in RNA-based cancer prognosis, where the SFT model achieved 0.77 accuracy on TCGA pairwise progression-free survival comparisons (which patient lived longer without progression?). In NHANES binary 10-year mortality prediction, the SFT model placed first with 0.89 accuracy, ahead of Gemini 3 Pro which was the best model previously tested in Longevity Bench. The regression variant of the NHANES time-to-death task yielded the lowest MAE among all tested systems (51.0 months). Our SFT model also achieved 0.60 accuracy on GTEx pairwise age comparisons (which sample is older?), a task category in which most previously tested LLMs struggled to exceed random baseline.

These results indicate that structured fine-tuning on aging-related tasks can close the performance gap between smaller models and frontier systems of substantially greater capacity. The improvements are seen in three distinct biodata modalities, suggesting that the acquired knowledge generalizes across data types rather than being confined to a single domain.

### Epigenetic age prediction

Predicting chronological age from DNAm is the most established use case of the aging clock methodology. We evaluated whether an LLM fine-tuned on DNAm data presented as text-formatted CpG beta values could emulate this function and potentially outperform specialized models. The training data comprised complementary prompt formats, featuring either a fixed set of CpG sites curated from multiple public clocks, or different sets of CpG sites representing important features of individual clock panels (see **Supplementary File 1**). These prompt collections contain actual DNAm profiles with beta values specified as decimals or percentages. In addition to these datasets, Longevity-LLM received training on prompts that taught it the biological significance, genomic annotation, of individual CpG sites and their connection to other biological entities.

After SFT, Longevity-LLM achieved an MAE of 5.91 years on 1,438 holdout samples, demonstrating the ability to reliably detect the aging signal in raw DNAm data (**Figure 2A**). After subsequent RFT, the model reduced the MAE to 4.34 years, significantly outperforming the Horvath multi-tissue clock on the same holdout set (MAE 4.61, Figure 2B).

**Figure 2.**
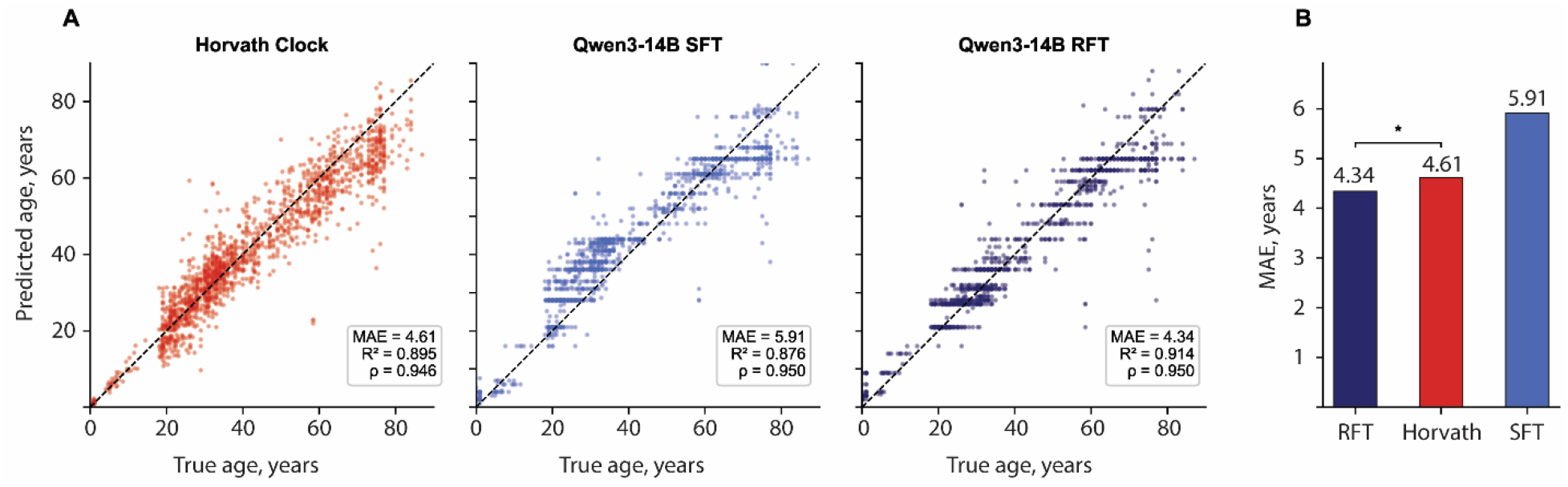
Longevity-LLM is a language model parsing the aging signal in DNAm profiles. (A) While SFT training on compiled datasets enables reliable age prediction, additional gains in performance can be achieved with RFT. Scatter plots show model performance in the holdout set of 1,438 DNAm samples. (B) The post-RFT Longevity-LLM significantly outperforms Horvath’s aging clock (two-sample bootstrap, p < 0.05). Dashed lines represent the unit line; ρ is Spearman correlation.

This result constitutes a proof-of-concept for a language model internalizing functionality previously delegated to purpose-built specialist tools. It is especially notable that Longevity-LLM features no architectural augmentations for DNAm data processing, such as dedicated prediction heads, special tokens, or separate treatment of numeric values. The base Qwen3 architecture, originally designed to process natural text, has proven fully sufficient for a task that previously required a dedicated statistical model.

### Proteomic age prediction and profile generation

We next evaluated whether Longevity-LLM could learn age-related proteomic patterns. The proteomic data comprised Olink Explore 3072 plasma profiles from 172 subjects across three independent cohorts (see Methods). Performance in age regression was compared against three proteomic aging clocks published in [ref, ref].

On 94 holdout samples, Longevity-LLM achieved an MAE of 7.9 years (**Figure 3A–B**), significantly below baseline error. The model performed within the accuracy range of purpose-built proteomic predictors, trained on a much larger UK Biobank dataset. Similar to the DNAm age prediction, Longevity-LLM’s base architecture was not modified to assist proteomic data processing.

**Figure 3.**
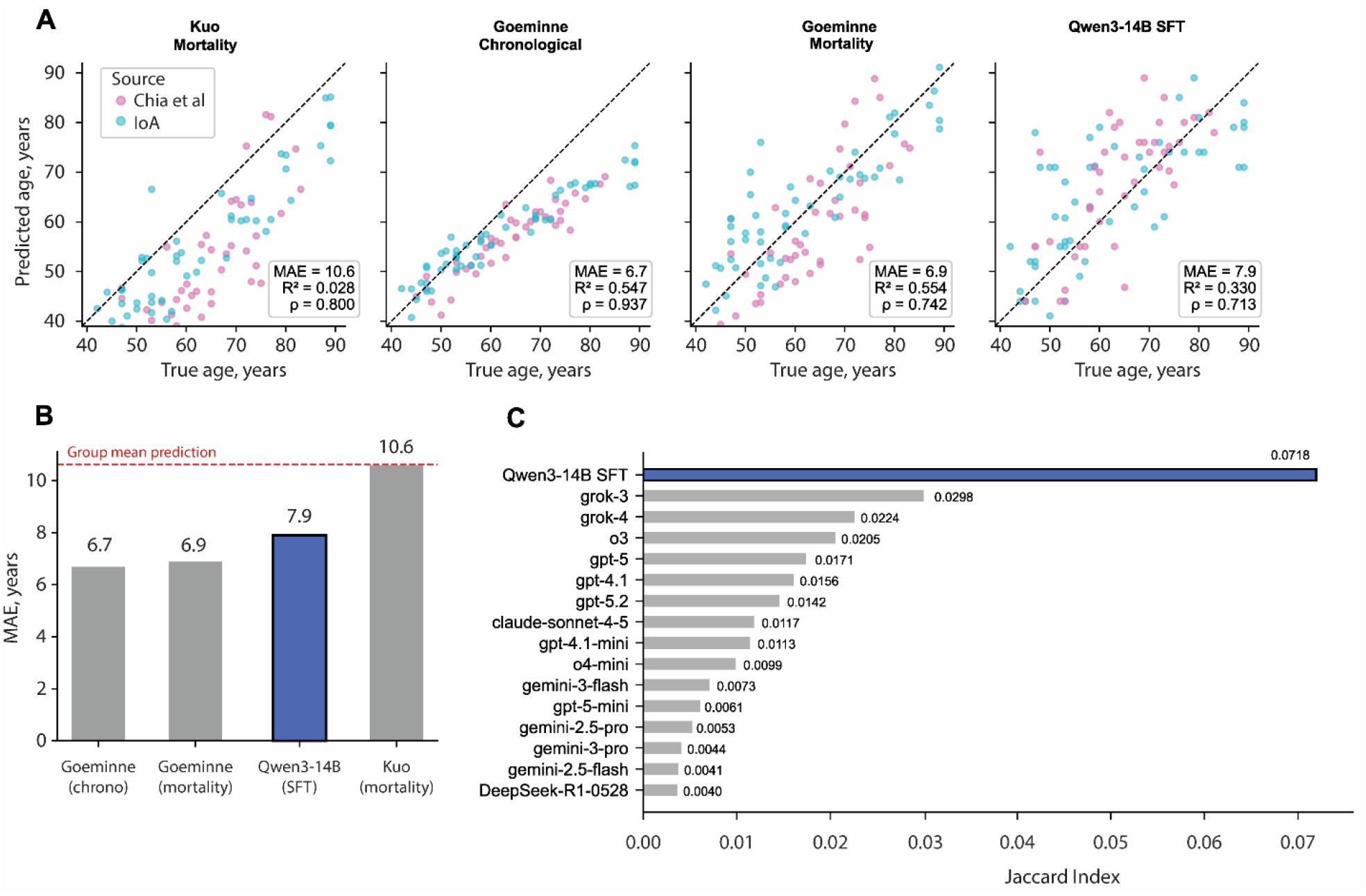
Longevity-LLM captures age-dependent signal in proteomic data. (A-B) Longevity-LLM’s age prediction accuracy falls within the range of specialist proteomic clocks. The test set contains 94 holdout samples from [ref, ref] ρ is Spearman correlation. (C) In proteomic profile generation, Longevity-LLM surpasses all tested frontier models, suggesting that it has internalized meaningful representations of proteome dynamics.

Beyond regression, we assessed Longevity-LLM’s capacity to generate biologically plausible proteomic profiles. When instructed to predict the 25 most abundant plasma proteins from a partial Olink profile conditioned on age and sex, the SFT model attained a Jaccard index of 0.072, which is significantly greater than any other tested frontier model (**Figure 3C**). This result indicates that fine-tuning yielded biologically meaningful representations of the aging plasma proteome, enabling the model to carry out tasks beyond numeric age mapping.

## Discussion

The experiments reported here demonstrate that aging clock distillation can produce a competitive multimodal aging model. The base model Longevity-LLM is based on, Qwen3-14B, was unable to generate valid predictions on any tested task, making the achievements in aging-related tasks even more impressive. After SFT and RFT, the resulting Longevity-LLM matched or exceeded frontier models in most Longevity Bench ^23^ tasks, while significantly outperforming the Horvath clock ^32^ in epigenetic age prediction. Moreover, Longevity-LLM demonstrated the ability to predict age from proteomic data, despite being trained with a much smaller dataset when compared to existing specialist clocks.

While conversational LLMs are typically considered a poor choice for the tasks encountered in omics domains, some teams propose novel architectures that account for highly-specific data formats and operations ^33^. Such models are not conversational and are hyper-focused on a narrow scope of tasks, which greatly limits their utility. For example, Evo2 is the state-of-the-art genomic foundation model, shown to generate gene sequences that carry out a predefined function ^34^. However, prompting it natural language is not possible, and the requests to the model need to be formatted as genomic sequence completion tasks. On a conceptual level, Longevity-LLM has shown that standard text transformers with no architectural modification still can serve as a valid development foundation for omics and aging research, maintaining the benefits of conversational models.

Aging clocks are currently the dominant analytical tool in aging research, which have long been seen as irreplaceable despite their many shortcomings ^13^. Most importantly, aging clocks are usually designed to support a single data modality and a fixed feature set. Compared to conventional clocks, Longevity-LLM can interact with any omics modality, partial observations, and its capabilities stretch beyond age or mortality regression. It represents a single access point to the multitude of views on aging biology captured in individual clocks, combining the statistical fidelity of dedicated models with the flexible instructiveness of LLMs.

The aging clock distillation we discuss in this work is but a particular instance of a more general approach. Similar prompt composition techniques and training regimens may be utilized to incorporate the knowledge embedded into virtually any type of a specialist model. This approach is actively leverage in the MMAI ecosystem to teach LLMs to navigate the broader interdisciplinary domain of biomedical research. As we show in our latest preprint, synthetic datasets and rewards constructed with them can lead to major improvements in a range of chemistry tasks related to retrosynthesis, pharmacokinetic property prediction, and toxicity profiling ^35^. Unlike chemistry, biology is not as tractable for defining objective reward function due to much greater system complexity. Biological systems are never isolated and extremely context-dependent, which leads to only limited success in biological and omics data synthesis. Despite these fundamental limitations, the sheer amount of omics data generated throughout the decades enables parsing the biological signal unobscured by confounder noise. The models of the Precious series ^36–39^, scGPT ^40^, Cell2Sentence ^41^, CpGPT ^2^ and MethylGPT ^3^ are foundation models capable of realistic omic data manipulation and generation for limited applications. However, much like genomic foundation models, they use custom architectures, tokenizers, and data preprocessing steps, which prevent their integration in the wider AI infrastructure. The distillation approach demonstrated herein is intended to remove the rift separating custom-built omics foundation models and state-of-the-art conversational LLMs, making it possible to query, and reason over factual biodata through natural language interfaces.

Our decision to build a monolithic aging model rather than a tool-based agent system was motivated by the goal of developing an internally consistent representation of molecular aging, so that it can be leveraged for purposes beyond those explored in this work. While agentic frameworks boast high accuracy in a wide array of tasks by the virtue of being an LLM-orchestrated ensemble of best-in-class models, biomedical research often requires internal consistency across biological scales ^42^. The ability to generalize between modalities and extrapolate partial information into actionable hypotheses is the key quality of research-grade AI. Such qualitative properties are unlikely to emerge in an ensemble simply by adding more independently developed tools. Moreover, each specialist model carries its own implicit assumptions about which features matter and why, while the orchestrating LLM has no means to reconcile them in a coherent way. A monolithic model, trained iteratively across modalities within shared parameters, creates the conditions for such cross-modal coherence. The proteomic generative results illustrate this principle, as Longevity-LLM outperformed frontier systems in generating protein profiles despite limited training data. This suggests that the biological information acquired from other domains may have contributed to better performance in the underrepresented modality.

This internal coherence, however, remains largely implicit in the current checkpoint and we aim to explore the cross-modal generalization hypothesis in follow-up studies. The predictive capabilities demonstrated here establish the foundation for the next stage of development: enabling Longevity-LLM to articulate the biological logic behind its predictions. Refining this capacity to reason from low-level molecular features to high-level phenotypic interpretation is the central objective of our next training iteration. Previously, we have demonstrated the power of biomedical AI systems by combining internally developed tools into a pipeline to nominate multiple drug candidates. For example in our idiopathic pulmonary fibrosis, we first identified TNIK as a “dual-purpose” target involved in both the pathology and general aging ^19,20,43^. Aging clock and hallmark of aging analytics contributed to its prioritization for clinical development ^44,45^. Then, our generative AI chemical platform Chemistry42 was employed to design and test *in silico* a range of small-molecule inhibitors ^46^. As preclinical models confirmed its beneficial properties ^44,47^, clinical trials began in earnest. Since then, the TNIK inhibitor rentosertib has passed phase-1 and phase-2 trials ^48–50^, as well as demonstrated geroprotector potential *in vitro* ^51^. The MMAI Gym initiative aims to combine the many separate AI models into one package capable of supporting and advancing a similar clinical program. The outlined distillation procedure is one component enabling minimal technological fragmentation in the upcoming drug development pipelines.

This work constitutes an interim report from the first sprint of our MMAI Gym for Science initiative. The model we ultimately aim to build would connect individual molecular measurements to pathway-level biology, interpret deviations from expected aging trajectories, and communicate its reasoning to researchers. To achieve this and circumvent the bottleneck of explicit reasoning dataset curation, we plan to intensify the RFT component of our project.

Incorporating more training datasets, including more test tasks, and scaling up RFT will be the top priorities in the Longevity-LLM v0.2 release. Altogether, our goal is to develop a model to serve as a companion in aging research rather than a prediction endpoint. Longevity-LLM v0.1 thus marks the beginning of a new stage in biogerontology by uniting disparate statistical models into a single system with promising growth potential.

## Abbreviations

CpG: cytosine-phosphate-guanine dinucleotide
DNAm: DNA methylation
GTEx: Genotype-Tissue Expression project
LLM: large language model
MAE: mean absolute error
MMAI: Multi-Modal AI Gym for Science
NHANES: National Health and Nutrition Examination Survey
R^2^: coefficient of determination
RFT: reinforcement fine-tuning
SFT: supervised fine-tuning
TCGA: The Cancer Genome Atlas
ρ: Spearman rank correlation coefficient

## Competing interests

All authors are employees of Insilico Medicine, a publicly traded drug development company (HKEX:3696.HK) developing AI applications for target discovery and drug design.

## Notes

https://insilico.com/MMAI

