## Supplementary File 1 for "The End of Aging Clocks: Training Foundation Models to Reason in Aging and Longevity"

### Training Prompt Schemas

This document provides representative prompt schemas for all training tasks used in training Longevity-LLM v0.1. Each schema shows the structural template with representative content; field values (biomarker names, units, gene annotations) were drawn from the actual training data. Tasks are grouped by data modality. Where multiple prompt formats exist for the same underlying task, the differences are noted.

All prompts were formatted for the Qwen chat template (<|im_start|>system, <|im_start|>user, <|im_start|>assistant). Multi-turn prompts contain six user–assistant exchanges per conversation, where the first five turns serve as few-shot demonstrations and the sixth is the target for prediction. Single-turn prompts contain one user turn and one assistant turn, with the target completion including a chain-of-thought reasoning trace in <think> blocks. Targets are enclosed in <answer> tags across all tasks.

### 1. General Population Survival (NHANES)

Two principal prompt formats were prepared for NHANES tasks. The full-profile format uses plain-text headers, fixed biomarker names, narrow unit conventions, and includes demographic, biometric, blood, medication, and questionnaire features. The blood-only format uses XML tags (<demographic>, <biometric>, <blood>), randomized biomarker synonyms and unit representations, and is restricted to blood biomarkers. A subset of blood-only prompts includes chain-of-thought reasoning traces in the target, in which the model reasons about individual biomarker values, identifies risk and protective markers, and states its conclusion before the final answer. These chains were generated programmatically and follow a pre-determined template pattern.

#### 1.1 Mortality prediction — binary

**Full-profile format:** train 10,000 (5.0M tokens), test 4,176 (2.0M tokens)

<|im_start|>system

You are a helpful assistant in chemistry and biology. Your task is to predict whether

a participant survives 10 years based on their biomarkers. ...<|im_end|>

<|im_start|>user

10-year survival prediction from biomarker data. Provide your final answer as a single

letter (A or B) inside <answer></answer> tags.

Did this participant survive 10 years post-examination?

A. No B. Yes

Patient data:

Demographic: 53-year-old male with some college of non-Hispanic black ethnicity

rates health as good.

Biometric: BMI 42.2 kg/m2, height 179.4 cm, waist circumference 129.5 cm, body

weight 135.7 kg, forced expiratory volume 3274.5 mL, diastolic blood pressure 81.3 mm Hg,

systolic blood pressure 126.0 mm Hg, pulse 44.7 bpm.

Blood: Albumin (g/L) 38.00; Alkaline phosphatase (U/L) 83.00; Blood urea nitrogen

(mg/dL) 12.00; ...[TRUNCATED]... Vitamin B12 (pmol/L) 261.25.

Questionnaire: Rates own health as good. Family history of diabetes.

Medication: Metformin (taken for 3 years), Lisinopril (taken for 5 years).<|im_end|>

<|im_start|>assistant

<answer>B</answer><|im_end|>

[...five more user–assistant turns...]

**Blood-only format (XML-tagged)**: train 19,021 (5.6M tokens), test 8,976 (2.8M tokens)

<|im_start|>user

Did this participant survive 10 years post-examination?

A. No B. Yes

Patient data:

<demographic>79-year-old female.</demographic>

<blood>Mean cell volume (fL) 97.60; BUN (mg/dl) 9.00; Vitamin B12 (pmol/l) 118.82;

Eosinophils % (pct) 1.00; ...[TRUNCATED]... Creatinine (µmol/l) 76.40;

RBC count (x10e12/L) 4.89; BILIT (µmol/L) 13.68.</blood><|im_end|>

<|im_start|>assistant

<answer>B</answer><|im_end|>

These formats use randomized biomarker synonyms (e.g., "BUN" vs. "Blood urea nitrogen") and unit representations (e.g., "mg/dl" vs. "mg/dL").

#### 1.2 Mortality prediction — pairwise

**Full-profile format:** train 10,000 (9.5M tokens), test 3,890 (3.6M tokens)

<|im_start|>user

Pairwise survival comparison from biomarker data.

Which participant had longer survival after examination?

A. Participant A B. Participant B

Participant A data:

Demographic: 76-year-old male with less than 9th grade school education of

non-Hispanic black ethnicity.

Biometric: BMI 29.1 kg/m2, ...

Blood: Albumin (g/dL) 3.90; ...[TRUNCATED]...

Medication: Donepezil (taken for 4 years).

Questionnaire: Has medical history of arthritis; cancer.

Participant B data:

Demographic: 80-year-old male with college graduate or higher of non-Hispanic

white ethnicity.

Blood: Albumin (g/dL) 4.50; ...[TRUNCATED]...<|im_end|>

<|im_start|>assistant

<answer>A</answer><|im_end|>

**Blood-only format (XML-tagged)**: train 15,000 (8.0M tokens), test 7,771 (4.4M tokens)

<|im_start|>user

Which participant had longer survival after examination?

A. Participant A B. Participant B

Participant A data:

<demographic>82-year-old male.</demographic>

<blood>MCV (fl) 98.90; LDL (mg/dL) 137.00; ...[TRUNCATED]...</blood>

Participant B data:

<demographic>85-year-old female.</demographic>

<blood>Creat (umol/l) 70.72; ...[TRUNCATED]...</blood><|im_end|>

<|im_start|>assistant

<think>

Participant A is 82, B is 85 years old. Participant A has borderline elevated creat:

106.10 vs 70.72 umol/L — elevated creatinine suggests declining glomerular filtration rate.

Participant B has borderline elevated bun: 3.93 vs 7.14 mmol/L — elevated BUN points to

reduced renal clearance or increased protein catabolism. Participant A has borderline

elevated wbc: 10.50 vs 6.90 x10e9/L — leukocytosis may indicate acute infection, chronic

inflammation, or a stress response. Participant A has 1 risk / 0 protective markers;

B has 0 risk / 0 protective markers. I have all the information that I need. My final

answer is:

</think>

<answer>B</answer><|im_end|>

The reasoning traces compare individual biomarker values between participants, tally risk and protective signals, and finalize with a conclusion.

#### 1.3 Time-to-event — categorical (multiclass)

**Full-profile format**: train 5,988 (3.2M tokens), test 1,318 (689K tokens)

<|im_start|>user

How long did this participant survive post-examination?

A. 15+ years B. 10-15 years C. 5-10 years D. 0-5 years

Patient data:

Demographic: 80-year-old male with less than 9th grade of school education.

Biometric: BMI 24.2 kg/m2, ...

Blood: Albumin (g/dL) 4.30; ...[TRUNCATED]...

Medication: Beclomethasone (taken for 2 months), ...[TRUNCATED]...

Questionnaire: Rates own health as good. Has medical history of angina pectoris;

stroke; current asthma.<|im_end|>

<|im_start|>assistant

<answer>D</answer><|im_end|>

**Blood-only format (XML-tagged):** train 7,288 (2.3M tokens), test 2,828 (912K tokens): Same categorical bins (0–5, 5–10, 10–15, 15+ years), XML-tagged blood-only features, randomized option order, multi-turn format with optional reasoning traces.

#### 1.4 Time-to-event — regression

**Full-profile format**: train 5,988 (3.1M tokens), test 1,318 (653K tokens)

<|im_start|>user

How many months did this participant survive post-examination?

Patient data:

Demographic: 73-year-old female...

Blood: Albumin (g/dL) 4.10; ...[TRUNCATED]...<|im_end|>

<|im_start|>assistant

<answer>156</answer><|im_end|>

**Blood-only format (XML-tagged):** train 5,096 (1.5M tokens), test 2,642 (802K tokens): Same task, XML tags, blood-only features, bare numeric target (months).

#### 1.5 Chronological age prediction — regression

**XML-tagged**: train 40,000 (11.2M tokens), test 8,976 (2.7M tokens)

<|im_start|>user

How old is this participant? Answer in years.

<demographic>Female.</demographic>

<biometric>BMI 47.8 kg/m2.</biometric>

<blood>γ-glutamyltransferase (U/L) 20.00; Insulin (uIU/ml) 15.63;

Segmented neutrophils percent (%) 18.50; ...[TRUNCATED]...</blood><|im_end|>

<|im_start|>assistant

<answer>32</answer><|im_end|>

**XML-tagged, with reasoning**: train 40,000 (11.4M tokens), test 8,976 (2.8M tokens): Same structure but with "Patient data:" header and a reasoning trace:

<|im_start|>assistant

<think>

Male participant with BMI 28.3 (overweight). HbA1c at 7.3 pct (elevated): HbA1c drifts

upward with age as glucose homeostasis deteriorates. Albumin at 31.0 gm/L (borderline low):

albumin declines with age due to reduced hepatic reserve and chronic inflammation.

Overall, 10 markers lean older and 4 lean younger.

</think>

<answer>69</answer><|im_end|>

The augmented format includes ×5 augmentation per participant (randomized synonym and unit selection); the reasoning format uses a fixed prompt set without re-augmentation.

#### 1.6 Age group classification — multiclass

Train 40,000 (12.2M tokens), test 8,976 (2.9M tokens)

<|im_start|>user

What is this participant's age group?

A. 18-30 B. 30-45 C. 45-60 D. 60-75 E. 75+

Patient data:

<demographic>Male.</demographic>

<biometric>BMI 25.9 kg/m2.</biometric>

<blood>RBC (x10e6/mcL) 5.04; RDW (%) 12.70; ...[TRUNCATED]...</blood><|im_end|>

<|im_start|>assistant

<answer>C</answer><|im_end|>

#### 1.7 Age comparison — pairwise

Train 15,000 (8.1M tokens), test 4,488 (2.7M tokens)

```
    <|im_start|>user
    Which participant is older, based on their clinical blood test results? Answer with
    one letter.
    A. Participant A  B. Participant B
        Participant A data:
    <demographic>Female.</demographic>
    <biometric>BMI 24.1 kg/m2.</biometric>
    <blood>CRP (mg/L) 1.20; ...[TRUNCATED]...</blood>
    Participant B data:
    <demographic>Male.</demographic>
    <blood>A1C (%) 5.60; ...[TRUNCATED]...</blood><|im_end|>
    <|im_start|>assistant
    <answer>B</answer><|im_end|>
```

### 2. Cancer Survival (TCGA)

**Pairwise**: train 9,000 (2.4M tokens), test 2,966 (806K tokens)

```
    <|im_start|>user
    Cancer progression-free survival prediction from clinical data.
    Which of the following two Prostate Adenocarcinoma cases has had a longer
    progression-free interval after the initial RNAseq screening:
        Patient-A: A 62-year-old male diagnosed with Stage IIIA tumor (TT3a, NN0, MM0);
    Patient-B: A 55-year-old male diagnosed with Stage IIA tumor (TT2a, NN0, MM0),
    Gleason score 6.
    Patient-A shows high activity in "Formation Of The Cornified Envelope" pathway
    (NES: 0.44, genes: KRT19, DSG2, ...[TRUNCATED]...).
    ...[up to 10 contrastive pathways]...<|im_end|>
    <|im_start|>assistant
    <answer>B</answer><|im_end|>
```

Prompts present pairs of patients with the same tumor type, supplemented with contrastive single-sample GSEA (ssGSEA) pathway annotations from Reactome. Multi-turn format.

### 3. Multi-Mutant Lifespan (Synergy)

Three prompt variants were prepared: regression (direct % prediction), full (QA with GO annotations and experimental context), and minimal (QA with gene names only). Regression: train 1,509 (687K tokens), test 64 (34K tokens). Full: train 1,509 (698K tokens), test 64 (35K tokens). Minimal: train 1,509 (234K tokens), test 64 (9K tokens).

**Regression**

```
    <|im_start|>user
    What is the lifespan effect of Cat(IF);Sod1(IF) compared to wild type?
        Organism: Drosophila melanogaster
    Genes: Cat: Cat (Catalase) | Sod1: Sod1 (Superoxide dismutase 1)
    GO: Cat: Function: metal ion binding, oxidoreductase activity, ...[TRUNCATED]...
    Experimental context: Temperature: 25°C Diet: cornmeal-agar medium ...[TRUNCATED]...
    Known effects: Cat(IF): -14.2% lifespan (39 days vs 45 days WT) |
    Sod1(IF): +27.8% lifespan (60 days vs 47 days WT)<|im_end|>
    <|im_start|>assistant
    <think>
    </think>
    <answer>+22.2%</answer><|im_end|>
```

These prompts present known single-mutant lifespan effects with GO annotations and experimental context, and ask for the combined double-mutant lifespan change.

### 4. Aging Clock Feature Importance

**Pairwise CpG comparison**: train 6,461 (922K tokens), test 817 (116K tokens)

<|im_start|>user

In the context of healthspan and longevity, methylation in which site serves as a more

definitive indicator of systemic physiological decline and mortality risk?:

A. cg23818978 or B. cg06533629?

cg23818978 is located at the Body of AMICA1.

cg06533629 is located at the TSS1500 of KLF14.

cg23818978 is located in the Open Sea, outside of any CpG island context.

cg06533629 is located in a CpG island./think<|im_end|>

<|im_start|>assistant

<think>

</think>

<answer>B</answer><|im_end|>

Pairwise comparison of two CpG sites with genomic annotations (gene location, CpG island context). The model selects which site is a stronger aging indicator based on published clock coefficients after correction for feature variance.

### 5. Transcriptomic Age Comparison (GTEx)

**Pairwise expression direction**: train 70,523 (9.8M tokens), test 10,000 (1.4M tokens)

```
    <|im_start|>user
    <question>In Skin (gender=M), comparing age 30.0-40.0 vs 19.95-30.0, what happens to CPA1 RNA expression with age?</question>
    <site>cg16847637: CPA1_1stExon; CPA1_5'UTR</site>
    <annotation>CPA1: linked via DNAm clock GrimAgeV1_simplified;
    CpG weight=37585.63385</annotation>
    <options>
        A. Expression increases
        B. Expression decreases
        C. No consistent change
    </options><|im_end|>
    <|im_start|>assistant
    <answer>A</answer><|im_end|>
```

These prompts ask whether a gene's expression increases or decreases with age in a specific tissue and sex, anchored by an associated CpG site and its clock weight.

### 6. DNA Methylation Age Prediction

#### 6.1 Multi-clock curated CpG panel

**Age regression from fixed CpG set**: train 5,076 (14.7M tokens), test 1,438 (4.2M tokens)

<|im_start|>user

Predict age from these cpg sites methylation values:

cg18181703: 0.47

cg17501210: 0.56

cg06419846: 0.78

...[353 CpG sites with decimal beta values]...<|im_end|>

<|im_start|>assistant

<answer>19.0</answer><|im_end|>

Uses a fixed panel of 353 CpG sites selected as the most widely represented and important features among DNAm clocks available through Biolearn, with raw beta values (0–1 scale). No gene annotations. Multi-turn few-shot format.

#### 6.2 Clock-derived CpG panels with annotations

**Annotated panels (reduced CpG sets with reasoning traces):** train 237,111 (207.1M tokens), test 59,835 (47.8M tokens)

```
    <|im_start|>user
    Predict the chronological age from DNA methylation data.
    Input: DNA methylation levels (%) for 150 CpG sites:
      cg20063906: 13%
      cg10699171: 69%
      ...[28–687 CpG sites with percentage values]...<|im_end|>
    <|im_start|>assistant
    <think>
    I will predict chronological age from DNA methylation data. The input contains
    methylation levels at 150 specific CpG sites, expressed as percentages.
    1. CpG site: cg20063906
       Methylation level: 13%
       Location relative to gene: Intergenic (chr11:85,929,769)
       Genomic region: Intergenic
       Epigenetic aging mechanism: This CpG is located in an intergenic region. The
       hypomethylation in this region is generally associated with potential influence on
       distal regulatory elements.

    2. CpG site: cg10699171
       Methylation level: 69%
       Location relative to gene: Intergenic (chr6:24,936,965)
       ...
    ...[per-CpG annotations for all sites: genomic coordinates, gene name, function,
    GO terms, KEGG/Reactome pathways, transcription factor regulators]...
    - Key genes associated with top 10 CpGs: ARID5A, SBF1, HCG18, MPPED1, TNFSF10,
      LPXN, PPP1R16B, TBR1
    </think>
    <answer>53</answer><|im_end|>
```

CpG sites drawn from epigenetic aging clock panels pulled from Biolearn. Methylation values reported as percentages. The reasoning trace provides rich per-CpG genomic annotations generated by an automated pipeline that performs GFF3-based gene lookup, CpG island classification, ENCODE cCRE mapping, and JASPAR TFBS scanning. Single-turn format.

**Full panels (larger CpG sets without reasoning)**: train 38,595 (121.9M tokens), test 9,574 (30.9M tokens)

```
    <|im_start|>user
    Predict the chronological age from DNA methylation data.
    Input: DNA methylation levels (%) for 1089 CpG sites:
      cg09773473: 88%
      cg07800658: 69%
      ...[92–1089 CpG sites with percentage values]...<|im_end|>
    <|im_start|>assistant
    <think>
    </think>
    <answer>62.0</answer><|im_end|>
```

Same task as above but with larger CpG panels (full clock feature sets) and empty reasoning traces. The two formats use complementary CpG site selections from the same set of aging clocks.

### 7. Transcriptomic Age Prediction

**Age regression from gene expression:** train 9,987 (29.2M tokens), test 2,466 (7.2M tokens)

<|im_start|>user

Predict age from these gene expression values: (Tissue: Heart)

ABCG1: 3.23

ABHD3: 5.46

ACAA1: 34.48

ACACB: 48.85

...[~300–500 genes with numeric expression values]...<|im_end|>

<|im_start|>assistant

<answer>68.0</answer><|im_end|>

Uses tissue-specific gene expression profiles with numeric intensity values. No gene annotations. Multi-turn few-shot format.

### 8. Proteomic Age Prediction

All proteomic tasks use Olink Explore 3072 proximity extension assay plasma proteomics data with NPX values. Multi-turn few-shot format.

#### 8.1 Binary age classification

Train 372 (571K tokens), test 94 (144K tokens)

<|im_start|>user

<platform>Olink Explore 3072 proximity extension assay (PEA) plasma proteomics.

Values are NPX (Normalized Protein eXpression, log2 scale). Top 200 consensus aging

proteins selected by cross-clock frequency analysis.</platform>

<question>Based on the plasma proteome profile of this male individual, what is their

age group?</question>

<olink>Top-100 most abundant proteins (NPX): GLO1 (5.67); BANK1 (5.07);

...[TRUNCATED]...</olink>

<options>A. 18-22 B. 55-65</options><|im_end|>

<|im_start|>assistant

<answer>B</answer><|im_end|>

#### 8.2 Pairwise age comparison

Train 5,000 (4.2M tokens), test 1,274 (1.1M tokens)

<|im_start|>user

<platform>Olink Explore 3072 proximity extension assay (PEA) plasma proteomics.

Values are NPX (Normalized Protein eXpression, log2 scale).</platform>

<question>Based on their plasma proteome profiles, which individual is older?</question>

<options>A. Patient-A B. Patient-B</options>

<olink>Top-100 differentially abundant proteins (NPX):

CASP1 (A: -8.89, B: -0.74); FOLR3 (A: 3.61, B: -4.33);

...[TRUNCATED]...</olink><|im_end|>

<|im_start|>assistant

<answer>B</answer><|im_end|>

#### 8.3 Age regression

Train 372 (560K tokens), test 94 (141K tokens)

<|im_start|>user

<platform>Olink Explore 3072 proximity extension assay (PEA) plasma proteomics.

Values are NPX. Top 200 consensus aging proteins selected by cross-clock frequency

analysis.</platform>

<question>What is this individual's exact age?</question>

<olink>GLO1 (5.70); KIF22 (4.63); ...[TRUNCATED]...</olink><|im_end|>

<|im_start|>assistant

<answer>58.0</answer><|im_end|>

#### 8.4 Generative protein prediction

Train 1,488 (5.2M tokens), test 376 (1.3M tokens)

<|im_start|>user

<platform>Olink Explore 3072 proximity extension assay (PEA) plasma proteomics.

Values are NPX. Neurology panel: Proteins pertinent to neurological diseases...

729 measured proteins: AAMDC;ABCA2;...[TRUNCATED]...</platform>

<question>This plasma proteomics sample is from a 55-65 year old M. Given 25 of the

top-50 most abundant proteins (NPX), predict the remaining 25 (in any order).</question>

<olink>GLO1 (5.70); KIF22 (4.63); RANBP1 (4.17); ...[25 proteins]...</olink><|im_end|>

<|im_start|>assistant

<answer>CSRP3;MYH9;TNNI3;...[25 predicted proteins]...</answer><|im_end|>

These prompts list all measurable proteins in the relevant Olink sub-panel, presents 25 of the top-50 most abundant, and instructs the model to predict the remaining 25.

### 9. Clock Coefficient Approximation

**Multi-omics biomarker importance comparison**: train 109,011 (101.3M tokens), test 21,564 (17.0M tokens)

```
    <|im_start|>user
    Increased methylation in blood at which CpG locus more strongly suggests a younger biological age?
A. cg02802055 or B. cg01131735.
Answer only A or B.
    cg02802055 maps to the Body region of C9orf78, in the north shore of a CpG island.
    cg01131735 maps to the TSS1500 region of FKBP10, in a CpG island.
    Gene: C9orf78 (chromosome 9 open reading frame 78). Summary: Located in cytosol and
    nucleoplasm. Key molecular functions include protein binding, RNA binding, and U5 snRNA
    binding. ...[extended gene annotation with GO, pathways, literature]...
    Gene: FKBP10 (FKBP prolyl isomerase 10). Overview: The protein encoded by this gene
    belongs to the FKBP-type peptidyl-prolyl cis/trans isomerase (PPIase) family.
    ...[extended gene annotation]...<|im_end|>
    <|im_start|>assistant
    <answer>A</answer><|im_end|>
```

Each conversation contains six pairwise biomarker comparisons spanning three omics modalities: CpG methylation (~70% of questions), transcriptomic gene expression (~20%), and plasma protein levels (~10%). Questions cover diverse aging-related traits including chronological age, biological age, mortality risk, cell proliferation history, gestational age, body composition, and smoking exposure. Each biomarker is accompanied by detailed gene annotations (function summary, GO terms, KEGG and Reactome pathways, transcription factor regulators, and literature findings). Ground-truth labels are derived from published aging clock coefficient magnitudes after feature variance corrections.
